# Lens Transmittance Shapes UV Sensitivity in the Eyes of Frogs from Diverse Ecological and Phylogenetic Backgrounds

**DOI:** 10.1101/783811

**Authors:** Carola A. M. Yovanovich, Michele E. R. Pierotti, Almut Kelber, Gabriel Jorgewich-Cohen, Roberto Ibáñez, Taran Grant

**Affiliations:** Department of Zoology, Institute of Biosciences, University of São Paulo, São Paulo, Brazil; Smithsonian Tropical Research Institute, Panama City, Panama; Department of Biology, Lund University, Lund, Sweden

**Keywords:** Anura, Diel pattern, Ocular media transmittance, Ultraviolet sensitivity, Vision, Visual ecology

## Abstract

The amount of short wavelength (UV, violet and blue) light that reaches the retina depends on the transmittance properties of the ocular media, especially the lens, and varies greatly across species in all vertebrate groups studied previously. We measured the lens transmittance in 32 anuran amphibians with different habits, geographic distributions, and phylogenetic positions and used them together with eye size and pupil shape to evaluate the relationship with diel activity pattern, elevation and latitude. We found an unusually high lens UV transmittance in the most basal species, and a range that extends into the visible spectrum for the rest of the sample, with lenses even absorbing violet light in some diurnal species. However, other diurnal frogs had lenses that transmit UV light like the nocturnal species. This unclear pattern in the segregation of ocular media transmittance and diel activity is shared with other vertebrates and is consistent with the absence of significant correlations in our statistical analyses. Although we did not detect a significant phylogenetic effect, closely related species tend to have similar transmittances, irrespective of whether they share the same diel pattern or not, suggesting that ocular media transmittance properties might be related to phylogeny.

## BACKGROUND

The lenses of animal eyes need to be transparent to allow the light they focus to reach its final destination, the photoreceptors in the retina. Among vertebrates, this condition is met invariably in the range ≈450–700 nm, but there is great variability across the UV-blue range of the spectrum (≈300–450 nm) [1]. Part of this variation is due to the chemical composition of the lens, as the proteins that comprise it absorb light ≤ 310 nm [1]. Thus, the bigger the lens, the greater the distance the light must traverse and the higher the probability that short wavelength light will be absorbed. Furthermore, light is also scattered by particles in the ocular media generally and the lens specifically, although this scattering is thought to be wavelength-independent [1]. A partial reduction in UV transmittance is, thus, a consequence of eye enlargement, which, in turn, enables high sensitivity and spatial resolution [2]. In addition to this trade off, there are optical benefits to modulating the spectral composition of light available for the retina, as short wavelengths are particularly prone to another type of scattering caused by small particles (Rayleigh scattering) and also to chromatic aberrations [1,3]. Furthermore, prolonged exposition to short wavelength light can cause photochemical damage to the retina [4], so it has been proposed that long-lived and diurnal animals might benefit from UV-absorbing lenses [5].

A widespread solution to the problems caused by short wavelength light is to add pigments to the lens to filter it. Such pigments have been spectrally and chemically characterized in some fishes [6,7], mammals [5], and the leopard frog [7] and inferred to exist in some birds [8]. A different strategy to cope with chromatic aberrations without filtering light are multifocal lenses, which allow light of different wavelengths to converge on the same focal plane *via* a specific refractive index gradient [9]. This mechanism obviously requires the whole lens to be exposed to the incoming light and would not work properly in eyes with round pupils that cover the periphery of the lens when contracted. Indeed, multifocal lenses strongly correlate with pupil shape in vertebrates: in a sample of 20 species from different tetrapod groups, all the species that have slit-shaped pupils also have multifocal lenses [10]. Thus, a combination of a highly short-wavelength transmissive lens with multifocal optics and a slit pupil can be an alternative to a pigmented, short-wavelength absorbing lens if photoreceptors have sensitivity peaks at short wavelengths or if light availability needs to be maximised, even at the cost of some scattering.

Lifestyle and geographic distribution determine the amount and spectral composition of light to which animals are exposed, both in the temporal (day, night) and spatial dimensions (latitude, elevation, and habitat type [11–13]), and, in turn, the lens (and occasionally the cornea) can selectively filter part of that light. Thus, it is reasonable to expect that lens transmittance properties would have evolved in such a way that they ‘match’ the light environment in which a given visual system performs. Accordingly, it has been hypothesised [14] that nocturnal animals would have highly transmissive lenses to maximise the number of photons that can reach the retina in a context in which they are scarce *per se*, and that diurnal animals for whom light is an ‘unlimited’ resource could afford to filter out part of the short-wavelength radiation to fine tune resolution while preventing the potential damage caused by the exposure to high amounts of that kind of radiation. Indeed, there seems to be a loose tendency for this to occur in fishes [6,15,16], snakes [17] and mammals [5],with exceptions in all cases, although the relationship between lens transmittance and diel pattern has not been statistically tested in any of them. Several studies have investigated the variability in lens transmittance at short wavelengths and its correlation with eye size, photoreceptor spectral sensitivity and a variety of natural history traits in fishes [16,18], lizards [19], snakes [17], birds [8,20], and mammals [5]. However, no broad comparative study of ocular media transmittance in amphibians has been pursued so far, and none of the studies in other lineages quantified those relationships in a formal phylogenetic context, so the interplay between ecology and evolutionary history in shaping the light transmittance properties in vertebrate eyes remains virtually unexplored.

In a previous study, we showed that the lenses of two species of anurans widely used as experimental models in vision research differ in more than 50 nm in the cut-off wavelength at which 50% of incoming light is transmitted (λ_T50_) [21]. Although that study also included another three closely related species, a broader sampling was needed to unveil the variability of lens transmittance across anuran species and lineages, and to assess potential relationships with their natural histories and with other properties of the visual system. In the present study, we assessed the lens transmittance, eye size, and pupil morphology of 37 species sampled from across the diversity of anurans and evaluated their relationship with the temporal and geographical environments that they inhabit.

## METHODS

### Sampling

We used eyes from 32 species of neobatrachian amphibians collected in their natural habitats in Brazil and Panama and one captive specimen of the basal species *Bombina orientalis* that died for reasons unrelated to the study (see Figure 1 for taxonomic distribution and phylogenetic relationships and electronic supplementary material S1A for details on identity and provenance of all specimens). We euthanized all the other animals by topical application of 2% benzocaine on the ventral skin buffered at pH 7 with sodium bicarbonate, until their breathing and cardiac activity ceased. In all cases, after death/euthanasia we enucleated the eye, freed the cornea by cutting along the *ora serrata*, cut through the vitreous to extract the lens, and removed the iris by cutting through the aqueous humour to obtain isolated corneas and lenses. All samples were freshly measured immediately after dissection.

**Figure 1.**
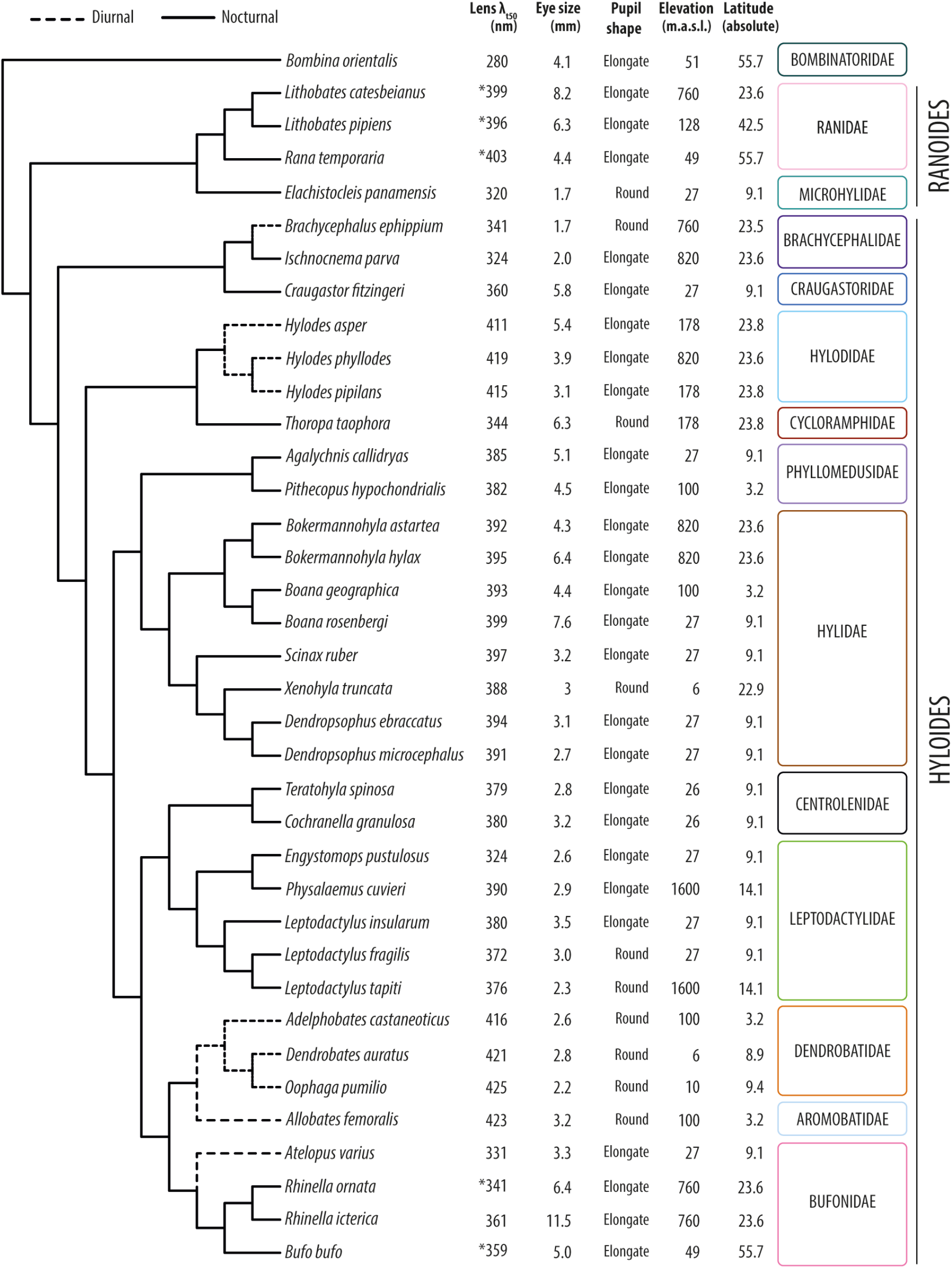
Summary of the values obtained for all the variables measured and compiled for the frog species included in this study in a phylogenetic context. (see electronic supplementary material S1F for the tree with branches lengths scaled to phylogenetic distances). All the lens λ_T50_ were calculated in this study except those marked with an asterisk (*), which were obtained from [21].

### Data collection

We measured lens transmittance (and corneal transmittance for some species) using the approach of Lind and co-workers [20], as follows. We placed the samples in a custom-made matte black plastic container with a circular fused silica window in the bottom and filled with phosphate-buffered saline (PBS). For small samples, black plastic discs with pinholes of 1 or 2 mm diameter were added on top of the silica window to ensure that all incoming light passed through the sample. We used an HPX-2000 Xenon lamp (Ocean Optics, Dunedin, FL) to illuminate the samples via a 50 μm light guide (Ocean Optics) and collected transmitted light using a 1000 μm guide connected to a Maya2000 spectroradiometer controlled by SpectraSuite v4.1 software (Ocean Optics). The guides were aligned with the container in a microbench system (LINOS, Munich, DE). The reference measurement was taken from the container filled with PBS. We smoothed the curves using an 11-point running average, and normalized to the highest value within the range 300–700 nm. From these data, we determined λ_T50_ as the wavelength at which the light transmittance was 50% of the maximum. The curves were cut for clarity in those cases in which the measurements at very low wavelengths were too noisy due to the low sensitivity of the spectrometer in that region of the spectrum.

We combined the lens transmittance data collected from the 32 species measured in this study with those from *Bufo bufo*, *Rhinella ornata*, *Lithobates catesbeianus*, *L. pipiens* and *Rana temporaria* that were available from a previous study [21], making a total of 37 species of 14 families. Given that corneal transmittance data were collected from just a handful of species, they were not included in the phylogenetic comparative analyses.

We used eye size compiled from descriptions of the species in the scientific literature as a proxy for lens optical path length. When these data were not available for a given species, we obtained them from colleagues or measured it from museum specimens (see electronic supplementary material S1B for the whole dataset of eye size values and sources, and S1C for validation of the method).

For pupil shape, we visually inspected photographs available online for each of the species and scored them as round or elongate (see electronic supplementary material S1D for details and thresholds on scoring criteria). Even though orientation (i.e. horizontally or vertically elongate) can have a differential effect on the sharpness of horizontal and vertical images [22] we did not distinguish between them because vertical slit pupils are extremely uncommon among anurans and would compromise the statistical analyses.

Diel activity pattern is somewhat labile in anurans and can vary for specific behaviours; however, most species are predominantly nocturnal (solid lines in Fig. 1) [23], with only a few lineages being predominantly diurnal, including the dendrobatoids (Aromobatidae + Dendrobatidae), hylodids, as well as *Atelopus* and *Brachycephalus* in our study (dashed lines in Figure 1) [23]. We opted to handle diel pattern as a binary variable, in line with previous work that uses this approach for different types of phylogenetic analyses [23]. Following this criterion, we scored *Scinax ruber* and *Lithobates pipiens*, which have been reported to be arrhythmic [23], as nocturnal based on our own fieldwork experience.

Given that elevation and latitude contribute to shaping light habitats, we also took them into account. We scored the elevation and latitude of the same specimens used to obtain the lens transmittance measurements, with two caveats. First, *Bombina orientalis* was captive bred in the pet trade in Lund, Sweden (elevation: 51 metres above sea level (m.a.s.l.), latitude: 55.7°), which is within the natural elevation of the species but is approximately 7° north of its northernmost distribution [24]. As such, we performed analyses both including and excluding this species. Second, Kennedy and Milkman [7] did not provide collection data for the *Lithobates pipiens* specimens they used to measure lens transmittance; given that the research was conducted at Harvard University, which is within the natural distribution of the species [25], we assumed they were collected nearby.

### Statistical analysis

We performed a phylogenetic comparative analysis to evaluate the linear relationships between lens λ_T50_, eye size, and pupil shape as predictor variables, and diel activity pattern, elevation, and latitude as response variables. Given the reports of a linear relationship between lens λ_T50_ and eye size in birds and mammals [5,8], we also tested this relationship explicitly. We used the phylogenetic hypothesis of Pyron [26] to control for the phylogenetic non-independence of the species in our sample (see electronic supplementary material S1E for details on how species missing in the tree were accommodated and 1F for the resulting tree).

We performed all analyses in R v3.6.0 using the packages *ape* v5.3 [27], *car* v3.0-3 [28], *GEIGER* v2.0.6.2 [29], and *nlme* v3.1-139 [30]. We used the function binaryPGLMM to run a phylogenetic generalized linear mixed model for binary data [31] for diel activity pattern.

For the continuous response variables of elevation and latitude, we performed phylogenetic general least squares analyses using the GLS function, correlation structures assuming a Brownian motion model of evolution, and transformation of the variance-covariance matrix of the phylogeny using Pagel’s *λ* values of 0, 0.01, 0.5, and 1 [32]. We tested for multicollinearity using variance inflation factors.

## RESULTS

The lens λ_T50_ of the 32 species measured in this study are spread throughout the UV-violet part of the spectrum, covering the range 280–425 nm (Figure 1), which also contains the values of the five species in our previous study [21]. The breadth of the range is similar in Hyloides and Ranoides, the two major lineages of neobatrachians that contain more than 90% of the anuran species diversity [33], although the upper boundary for the latter seems to be lower than for the former (Figure 1). The lowest value of the range (λ_T50_=280 nm) is that of *Bombina orientalis*, a comparatively basal species that is phylogenetically distant from the rest of the lineages in our sample (Figure 1, electronic supplementary material S1F).

We also measured the transmittance of the corneas in eight species from our sample. For most of them the λ_T50_ was within the range ≈320–345 nm, irrespective of the lens transmittance properties (Figure 2, electronic supplementary material S1G). However, *Physalaemus cuvieri* has a cornea λ_T50_=293 nm (Figure 2), and that is probably also the case for *Xenohyla truncata*, although the precise λ_T50_ value could not be calculated for the latter (electronic supplementary material S1G).

**Figure 2.**
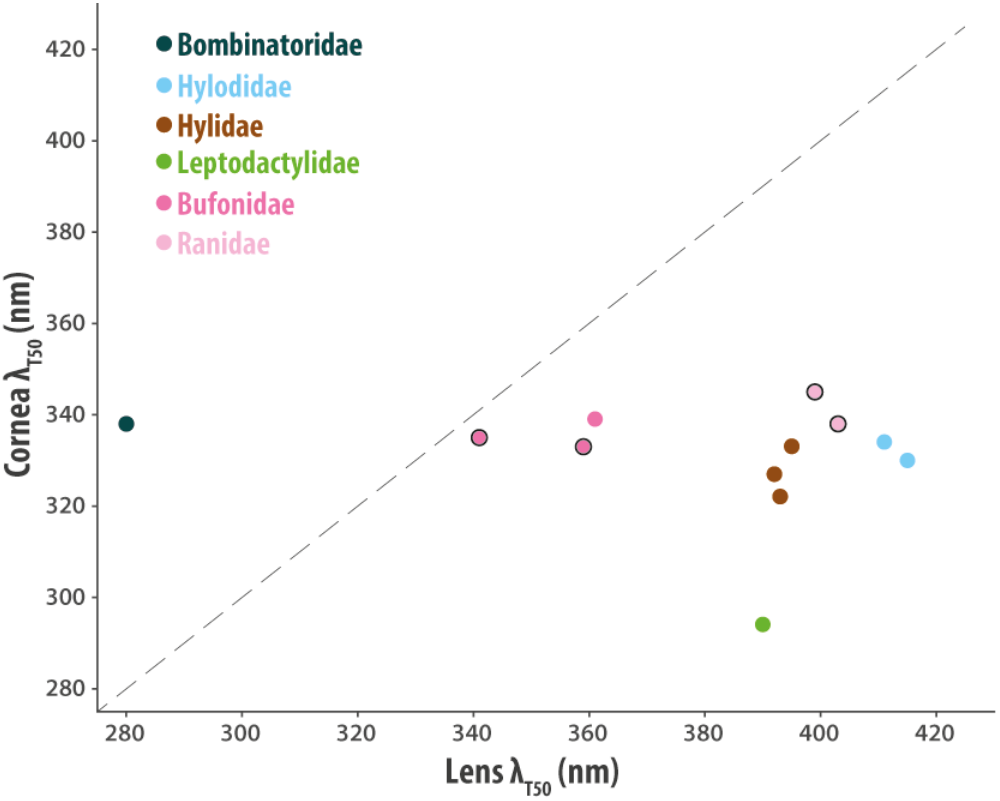
Relationship between cornea and lens transmittance for a subset of the species used in the study. Each data point represents one species, and the data for the points with a black outline were obtained from a previous study [21]. The diagonal defines the regions in which light transmittance of the eye as a whole is limited by the cornea (above the line) and by the lens (under the line). See electronic supplementary material S1G for corneal transmittance curves, λ_T50_ values and species identification.

The lens transmittance curves in our sample show the expected sigmoidal shape (Figure 3, electronic supplementary material S1H), with some variation in the slope of the short-wavelength cut-off and the saturation at long wavelengths. A noticeable feature in the shape of some of the curves is a localised increase in transmittance in the range ≈310–340 nm (*Hylodes phyllodes*, *Oophaga pumilio*, *Dendropsophus microcephalus* and *Cochranella granulosa* in Figure 3 and other species closely related to each of them; electronic supplementary material S1H), as well as the shoulder in the same wavelength range in *Craugastor fitzingeri* and *Brachycephalus ephippium* (Figure 3).

**Figure 3.**
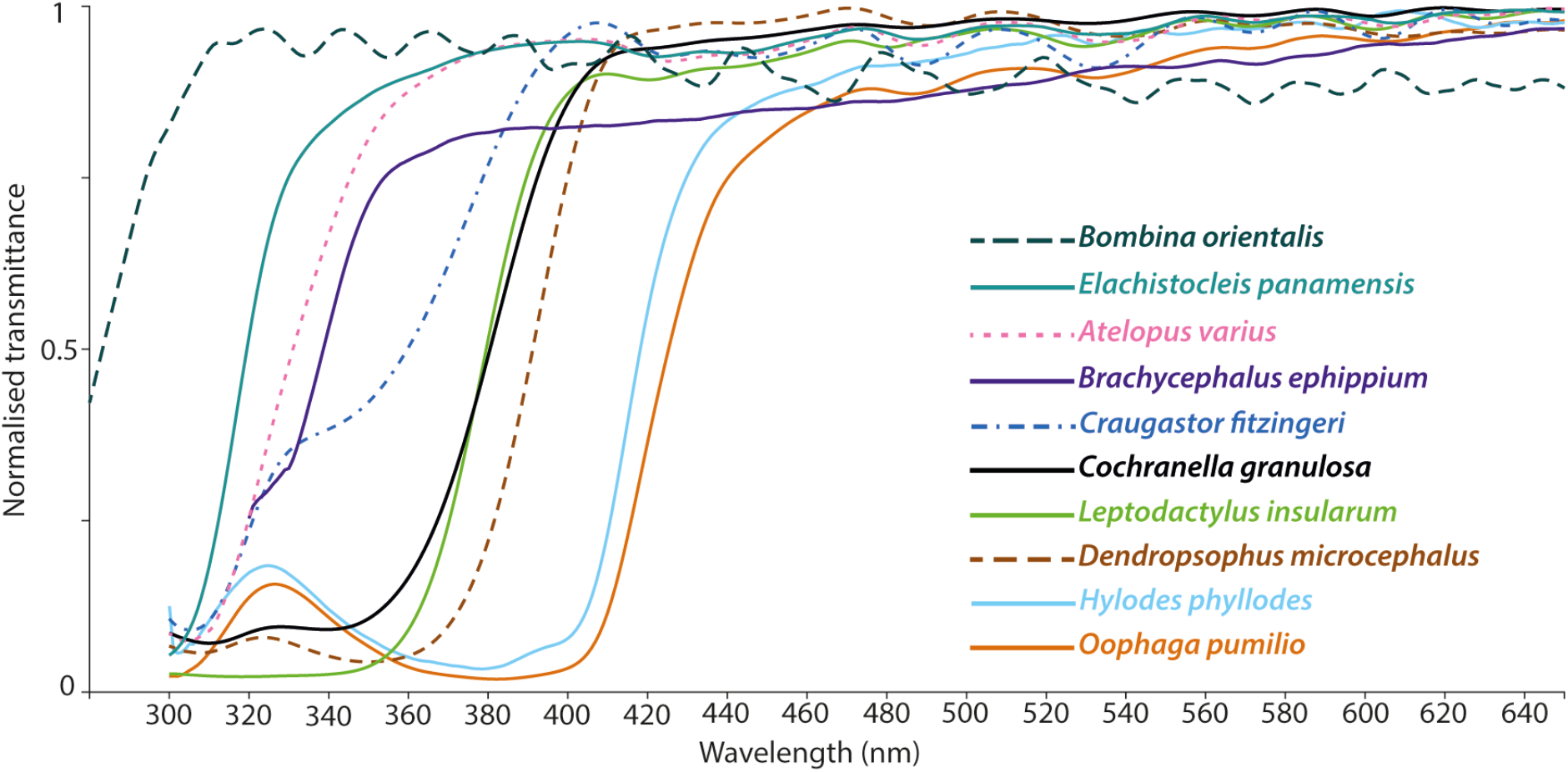
Lens transmittance curves from some of the species in the study. The X axis is cut at 650 nm to ease visualisation of the curves at short wavelengths. See electronic supplementary material S1H for curves from the rest of the species.

We found no correlation between lens λ_T50_ and eye size, either controlling (Pagel’s *λ* = 1; regression coefficient = 3.7164, p = 0.1777) or not controlling (Pagel’s *λ* = 0; regression coefficient = 0.4643, p = 0.868; Figure 4) for phylogeny.

**Figure 4.**
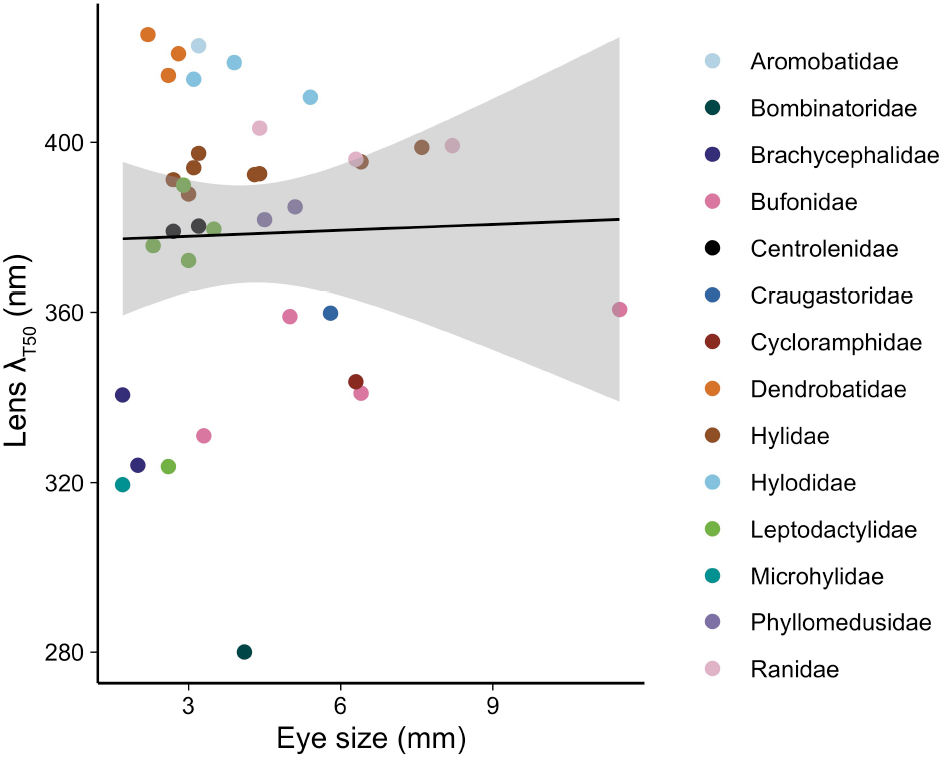
Linear relationship between lens transmittance and eye size. The shaded area is the 95% confidence interval. See electronic supplementary material S1J for results obtained with a subset of species bearing putatively un-pigmented lenses.

We also found no significant relationship between lens λ_T50_, eye size, and pupil shape (predictor variables) and diel pattern, elevation, and latitude (response variables) for any of the models (p > 0.09; see electronic supplementary material S1I).

## DISCUSSION

### Limits of UV transmittance in anuran eyes

Our results show that lens transmittance among anurans spans a range similar to that of other vertebrates such as fishes, snakes, lizards, birds and mammals [5,6,8,16–19]. Our sample covers virtually the whole latitudinal range of geographical distribution of the group, a broad altitudinal range (0–1600 m.a.s.l.), and includes diurnal and nocturnal species (Figure 1), thus covering the diversity of light spectral composition and illumination intensity to which most adult terrestrial anurans are exposed. It would be interesting to investigate whether the lenses of aquatic species and/or those that live in very high elevations show specific patterns within this range, or significant departures from it.

The lower limit in the range of lens λ_T50_ among the anurans in our sample is intriguing, and more than 20 nm shorter than the most extreme cases reported so far in vertebrates: the porcupine fish *Diodon hystrix* (301 nm, [6]), the sand-dwelling lizard *Calyptommatus nicterus* (303 nm, [34]), the African house snake *Boaedon (Lamprophis) olivaceus* (306 nm, [17]), and the Japanese quail *Coturnix japonica* (≈310 nm, [35]). However, the unusually high lens UV transmission in *B. orientalis* has no functional relevance in terms of light availability for the retina, since the cornea of this frog has a λ_T50_=338 nm (similar to other species with higher lens λ_T50_ values; Figure 2 and electronic supplementary material S1G). This means that the amount of ultraviolet light that can effectively reach the photoreceptors is comparable to that in frogs with lens λ_T50_≈335–340 nm. Thus, in this particular species the light transmittance of the eye as a whole is limited by the cornea rather than the lens (Figure 2), as is the case in quails [35]. However, this is likely the exception rather than the rule and not necessarily the case in other species in our sample with short lens λ_T50_ values, such as *Ischnocnema parva*, *Engystomops pustulosus* or *Elachistocleis panamensis*. Even though we do not have data on the corneal transmittances of any of them, the data from other species in our sample (e.g. *Physalaemus cuvieri*) show that anuran corneas can in some cases transmit virtually all light down to 300 nm and further. Furthermore, a broad sample of fishes showed a presumptive trend for corneas to be more transmissive than lenses for any given species [16], although this relationship has not been formally tested for any vertebrate group.

The remarkably low lens λ_T50_ that we found in *Bombina orientalis* is intriguing both from the point of view of lens structure and of the evolutionary history of UV transmittance in anurans. Biological tissues in general transmit only UV radiation >310 nm, as aromatic aminoacids absorb shorter wavelengths [1], so this lens might have either a specific spatial distribution of proteins, a low concentration of them, a particular crystalline type or a combination of all these factors to achieve a λ_T50_ value of 280 nm.

It is tempting to wonder whether this species is representative of the ancestral state of lens transmittance among anurans, given its basal phylogenetic position relative to the other species in our analyses and the fact that its cornea filters potentially damaging UV radiation, which might have relaxed selective pressure to make the lens less transmissive. Data from caudates —the sister group of anurans— seems to be limited to the salamander *Salamandra salamandra*, the newt *Cynops pyrrhogaster* and the axolotl *Ambystoma mexicanum* [1], all of which are deeply nested within Caudata [26]. Although lens λ_T50_ values have not been published for any of them, a gross visual estimation from the available curves (in figure 3b from reference [1]) suggests a range of 310–320 nm, which is intermediate between *Bombina* and the anuran species in our study. As our sample was too phylogenetically sparse to allow meaningful evaluation of the evolutionary history of this trait through ancestral state reconstruction, further studies focused on a richer sampling of anuran basal groups, as well as caudates, would be required to clarify this point.

### How is UV filtering achieved in anuran lenses?

Variation in the shape of the transmittance curves among lenses that absorb part of the UV radiation is quite a common theme in vertebrates, and in particular local increases at short wavelengths can be seen in lens transmittance curves from mammals [5] and snakes [17]. However, the only group in which this phenomenon has been thoroughly studied are fishes, for which several pigments drive these patterns [6]: fishes with curves like the ones for *Hylodes phyllodes* and *Oophaga pumilio* have a pigment with peak absorption at ≈370 nm, whereas other species of fishes with shoulders in their transmittance curves similar to those of *Brachycephalus ephippium* and *Craugastor fitzingeri* have two different pigments with peak absorptions at ≈320–330 and 360 nm. Finally, fish lenses with smooth curves and high λ_T50_ values like the one from *Leptodactylus insularum* in our sample have high concentrations of either the 360 nm pigment or both the 320–300 nm and 360 nm pigment [6]. Curves with very subtle local increases at short wavelengths similar to those of *Dendropsophus microcephalus* and *Cochranella granulosa* in our sample have not been reported in fishes, but are present in some mammals such as the meerkat, in whom lens pigments with absorption maxima at 360–370 nm have been extracted [5].

The similarity between fishes and anurans in the overall shape of transmittance curves for lenses of different species suggests that a number of pigments are involved in generating those patterns in the latter, as they are in the former. However, no comparative studies of lens pigmentation have been conducted in anurans. The only species for which a lens pigment has been extracted is the leopard frog *Lithobates pipiens*; its absorbance peaks at 345 nm and it was not identified [7]. However, this absorbance profile does not match any other pigment identified in the lenses of fishes or mammals [1], so it is very likely that its chemical identity is different.

The presumptive presence of pigments in some anuran lenses can explain the lack of correlation between lens transmittance and eye size in our analyses, as that relationship holds only for unpigmented lenses [1]. It is thus possible that a relationship between the two variables exists in amphibians, as it does in birds [8,35], mammals [5] and some fishes [36], but is masked by the pigmented lenses in our sample. Absence of lens pigments has been demonstrated for 33 species of fishes with smooth transmittance curves and lens λ_T50_≈310–340 nm [6]. Interestingly, if the linear regression for our sample is performed only with the six species that also have smooth transmittance curves and lens λ_T50_≈310–340 nm, the relationship between lens transmittance and eye size has an excellent fit (R^2^=0.96; electronic supplementary material S1J). If variation in the occurrence of pigment is confirmed for frog lenses, the relationship between lens transmittance and eye size should be re-tested among the species that fulfil the requirement of absence of pigment.

### Potential factors driving the relationship (or lack thereof) between lens transmittance and diel pattern

All studies that have qualitatively tested the hypothesis that lenses of diurnal and nocturnal vertebrates are UV-absorbing and UV-transmissive, respectively, mention deviations from this expected distribution pattern [5,15,17] that can seem anecdotal in each particular case, but taken together they always point in the same direction: all nocturnal species have UV-transmissive lenses and all species with UV-absorbing lenses are diurnal, but some diurnal species have UV-transmissive lenses (Figure 5).

**Figure 5.**
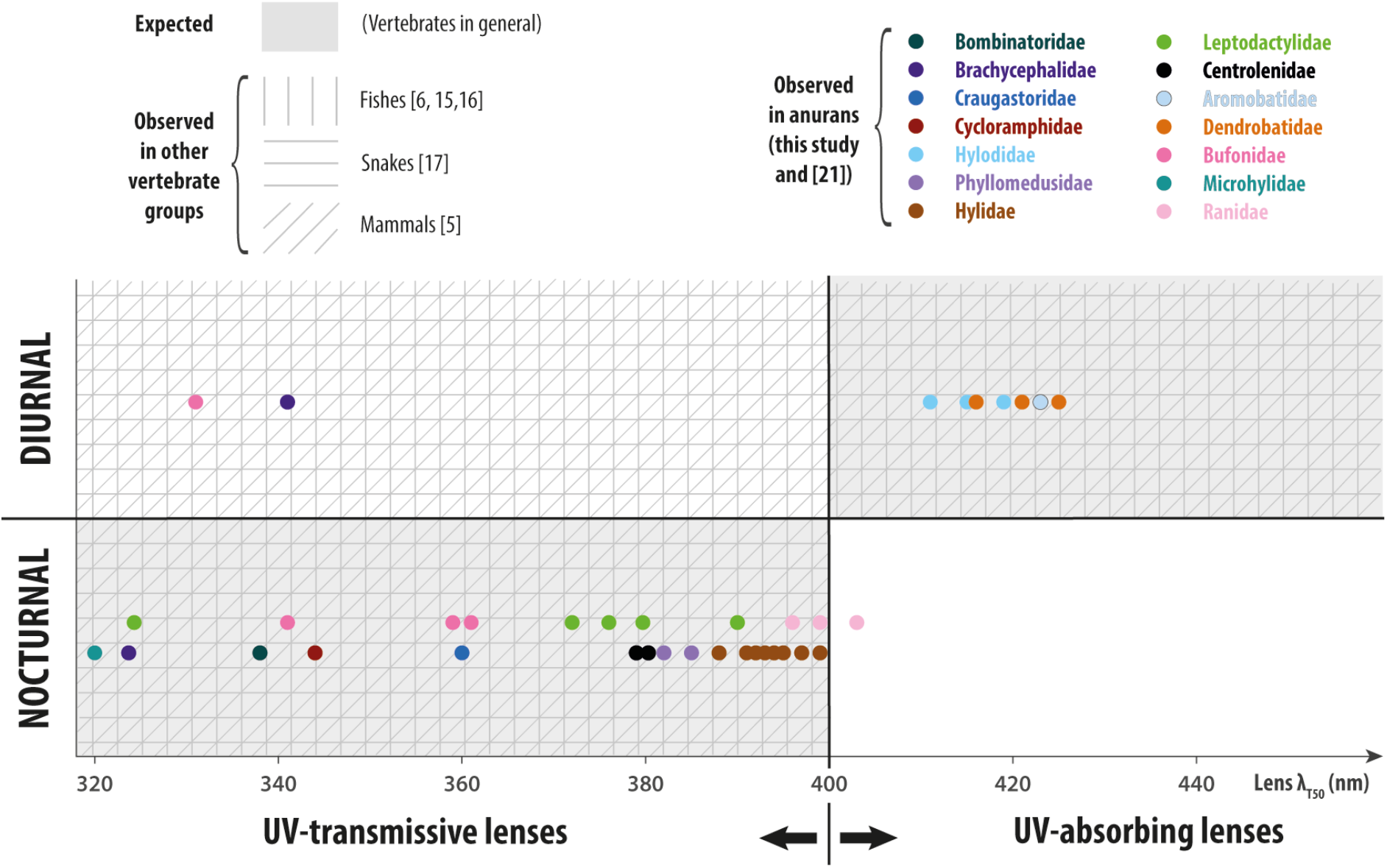
Distribution of lens transmittance values in diurnal and nocturnal representatives of different vertebrate groups. Each dot represents one anuran species. For Bombinatoridae (represented by *Bombina orientalis*), we show the cornea (rather than the lens) λ_T50_ as it is the limiting component for UV transmission in this species. The boundary between UV transmission and absorption is an artificial barrier within a continuum and is meant to aid grouping and visualization. A compilation of exact lens λ_T50_ values and diel patterns of species in groups other than anurans was outside the scope of this study, so those data are shown as general presence/absence patterns within each quadrant.

In this scenario, it comes as no surprise that there are diurnal anuran species in our sample on both sides of the UV transmission axis and no significant correlation of lens transmittance with diel pattern (and by extension, with the remaining variables that influence intensity and spectral composition of the light environment).

Despite their shared propensity to transmit at least part of the incoming UV radiation through their ocular media, the benefits might differ among nocturnal species from different vertebrate groups. Nocturnal vision in vertebrates is driven by rod photoreceptors, which typically have a peak spectral sensitivity outside the UV range at approximately 500 nm [37]. In addition, rods, as well as all other vertebrate photoreceptors, have a secondary lower, broader peak in the UV range (the β-band) [37] whose contribution to overall photon catch can become relevant and improve visual sensitivity in dim light when the total number of photons is extremely limited. In the case of amphibians, their rods are generally bigger— and thus more sensitive—than those of other vertebrates [38], so the contribution of UV light to overall visual sensitivity might not be as crucial as in other groups; indeed, the lenses of many nocturnal frogs are close to the boundary between UV-transmissive and UV- absorbing (e.g. some hylids and ranids, Figure 5) and absorb almost all light in the region of the β-band [21]. However, anurans and some caudates are unique among vertebrates in having a second rod type with peak sensitivity at ≈435 nm, in addition to the typical one at ≈500 nm [37,39]. This dual rod system allows frogs to retain the ability to discriminate colours down to light intensities in which other vertebrates become colour-blind [40,41], and its proper functioning might be relevant for many of the ≈80% of anuran species that are nocturnal [23]. In this context, it becomes crucial that the lens does not absorb too much short wavelength light; λ_T50_=403 nm already reduces a significant amount of the light that can reach the retina in *Rana temporaria* and removes almost completely the β-bands of both rods’ spectral sensitivity curves [21], and higher values could affect the sensitivity peak of the blue-sensitive rods, becoming detrimental to the performance of the visual system of nocturnal frogs in the dimly lit environments they inhabit.

As is the case with other vertebrates, there is no clear reason why some of the diurnal frogs in our sample depart from the expected UV-absorbing lenses. Filtering short-wavelength radiation can help reduce scattering and chromatic aberrations, thus improving spatial resolution, as has been suggested for animals that depend on sharp vision, such as raptors [8,20] and gliding snakes [17], and share UV-absorbing lenses. In the case of frogs, enhanced spatial resolution could be advantageous to species that use visual displays. All the diurnal representatives in our samples use them [42–45], suggesting that the optical problems caused by UV light are not serious enough—probably due to the small size of their eyes—to drive the selective pressure towards longer-wavelength shifted lens λ_T50_ values in all cases. An alternative explanation could be that, in some species, chromatic aberrations, if relevant, are dealt with by multifocal lenses rather by than UV-absorbing ones. Multifocality has only been tested in two anurans: the bufonid *Rhinella marina* (formerly *Bufo marinus*) has a multifocal lens, while the dendrobatid *Phyllobates bicolor* does not [10]. These data complement the differences in UV transmittance between the diurnal bufonid (*Atelopus*) and dendrobatids in our sample but are too limited to speculate about potential generalities. It would be interesting to obtain information about both lens transmittance and focal optics in the same species for a variety of anuran lineages, which would enable well-grounded hypotheses to be formulated about potential relationships between the two variables.

There is the possibility that UV light carries information valuable to some species in ways that are beyond both our knowledge of their visual ecology and our ability to imagine, given our own blindness in that part of the spectrum. In a recent study, it was shown that UV- and violet-light sensitivity can resolve habitat structure by increasing the contrast between the upper and lower surfaces of leaves to an extent that depends on the geometry of the canopy [46]. This previously unforeseen result showcases the way in which subtleties obscured by broad temporal and spatial habitat classifications (e.g. diurnal, nocturnal, open, or forest) can be the actual driving force underlying specific adaptations in traits that seem to deviate from expected patterns.

Finally, the ‘mismatch’ between lens transmittance and diel pattern in anurans in particular and in vertebrates in general might be related to phylogeny. For example, although we did not detect a significant phylogenetic effect in our data, it is evident that species of the same families tend to have similar lens transmittance properties, irrespective of whether they share the same diel pattern or not (e.g., Brachycephalidae, Bufonidae). This shows that within certain transmittance ranges and in the absence of highly specialised ecological demands, fluctuations in diel patterns within lineages have occurred without major departures from ancestral lens transmittance properties. The caveat that the phylogenetic constraints can be overridden by other factors is illustrated in our sample by the fact that the Túngara frog *Engystomops* (formerly *Physalaemus*) *pustulosus* has a lens λ_T50_ value approximately 50 nm shorter than its close relative *Physalaemus cuvieri* and all other leptodactylids in our sample (Figure 1). We hope that our work will encourage further research and data collection on ocular media transmittance from additional amphibian species to broaden our sampling, thus enabling robust testing of phylogenetic signals.

## ACKNOWLEDGEMENTS

We are thankful to Miguel Trefaut Rodrigues for providing specimens and to Marco A. de Sena, José Mario Gellere, Hugo Bonfim, Isabela R. Cavalcanti, Sergio M. de Souza, Agustín Camacho Guerrero, Délio Baêta, Ariadne F. Sabbag, Carla M. Lopes, Jhon Jairo Ospina Sarria, Andrés Brunetti, Hélio R. da Silva, Edivaldo Vasconcelos de Farias, Pedro Henrique Salamão Gananga, Alfredo Pedroso dos Santos, Síria Ribeiro, and Ricardo Cossio for help with logistics and fieldwork. Our gratefulness also goes to Thais Condez, Ariadne F. Sabbag, Mariane Targino, and Marco A. Rada García for generously sharing their data on eye sizes.

## ETHICS

The specimen of *Bombina orientalis* had been kept as a pet in Lund, Sweden and was donated to us on the day of its decease. Specimens from Brazil were collected and euthanized under licenses 13173-2 and 54599-3 from Chico Mendes Institute for Biodiversity Conservation and Biodiversity Authorisation and Information System (ICMBio/SISBIO). The method for euthanasia was selected and applied according to Resolution N° 37 of the Brazilian National Council for Control of Animal Experimentation. Specimens from Panama were collected under the permits SE/A-47-18, DAPB-0407-2019, SC/A-7-19, DAPB-0407-2019, SE/AP-8-19, issued by the Ministerio de Ambiente.

## DATA, CODE AND MATERIALS

Additional details about methods and results, as well as all transmittance datasets, are available in the Supplementary Material.

## AUTHORS’ CONTRIBUTIONS

CAMY, MERP, AK and TG conceptualized the study. CAMY, MERP, RI and TG conducted fieldwork. CAMY, MERP, GJC and RI collected data. CAMY, MERP, AK and TG analysed data. CAMY wrote the manuscript with feedback from all authors. All authors approved the final version of the manuscript.

