## supplementary material for "Lens Transmittance Shapes UV Sensitivity in the Eyes of Frogs from Diverse Ecological and Phylogenetic Backgrounds"

**Yovanovich CAM, Pierotti MER, Kelber A, Jorgewich-Cohen G, Ibáñez R & Grant T.**

### A. Specimens used for lens and cornea transmittance measurements

| Species | Sex | Source | Locality (State) | Country |
|---|---|---|---|---|
| *Bombina orientalis* | NR | CB | Lund | Sweden |
| *Brachycephalus ephippium* | F | FT | Mogi das Cruzes (SP) | Brazil |
| *Brachycephalus ephippium* | M | FT | Mogi das Cruzes (SP) | Brazil |
| *Ischnocnema parva* | M | FT | Boracéia (SP) | Brazil |
| *Craugastor fitzingeri* | M | FT | Gamboa | Panama |
| *Thoropa taophora* | F | FT | São Sebastião, (SP) | Brazil |
| *Thoropa taophora* | M | FT | São Sebastião (SP) | Brazil |
| *Hylodes asper* | M | FT | São Sebastião (SP) | Brazil |
| *Hylodes phyllodes* | M | FT | Boracéia (SP) | Brazil |
| *Hylodes pipilans* | M | FT | São Sebastião (SP) | Brazil |
| *Agalychnis callidryas* | M | FT | Gamboa | Panama |
| *Pithecopus hypochondrialis* | M | FT | Fazenda Treviso (PA) | Brazil |
| *Bokermannohyla astartea* | M | FT | Boracéia (SP) | Brazil |
| *Bokermannohyla astartea* | M | FT | Boracéia (SP) | Brazil |
| *Bokermannohyla hylax* | M | FT | Boracéia (SP) | Brazil |
| *Boana geographica* | F | FT | Fazenda Treviso (PA) | Brazil |
| *Boana geographica* | F | FT | Fazenda Treviso (PA) | Brazil |
| *Boana rosenbergi* | M | FT | Gamboa | Panama |
| *Boana rosenbergi* | M | FT | Gamboa | Panama |
| *Scinax ruber* | M | FT | Gamboa | Panama |
| *Scinax ruber* | M | FT | Gamboa | Panama |
| *Xenohyla truncata* | NR | FT | Maricá (RJ) | Brazil |

| Species | Sex | Source | Locality (State) | Country |
|---|---|---|---|---|
| *Dendropsophus ebraccatus* | M | FT | Gamboa | Panama |
| *Dendropsophus ebraccatus* | M | FT | Gamboa | Panama |
| *Dendropsophus microcephalus* | M | FT | Gamboa | Panama |
| *Dendropsophus microcephalus* | M | FT | Gamboa | Panama |
| *Engystomops pustulosus* | M | FT | Gamboa | Panama |
| *Engystomops pustulosus* | M | FT | Gamboa | Panama |
| *Physalaemus cuvieri* | NR | FT | Alto Paraiso de Goiás (GO) | Brazil |
| *Physalaemus cuvieri* | NR | FT | Alto Paraiso de Goiás (GO) | Brazil |
| *Physalaemus cuvieri* | NR | FT | Alto Paraiso de Goiás (GO) | Brazil |
| *Leptodactylus insularum* | M | FT | Gamboa | Panama |
| *Leptodactylus insularum* | M | FT | Gamboa | Panama |
| *Leptodactylus fragilis* | F | FT | Gamboa | Panama |
| *Leptodactylus fragilis* | F | FT | Gamboa | Panama |
| *Leptodactylus tapiti* | M | FT | Alto Paraiso de Goiás (GO) | Brazil |
| *Leptodactylus tapiti* | M | FT | Alto Paraiso de Goiás (GO) | Brazil |
| *Teratohyla spinosa* | M | FT | Soberania National Park | Panama |
| *Teratohyla spinosa* | NR | FT | Soberania National Park | Panama |
| *Cochranella granulosa* | M | FT | Soberania National Park | Panama |
| *Cochranella granulosa* | M | FT | Soberania National Park | Panama |
| *Allobates femoralis* | M | FT | Fazenda Treviso (PA) | Brazil |
| *Adelphobates castaneoticus* | M | FT | Fazenda Treviso (PA) | Brazil |
| *Dendrobates auratus* | M | FT | Ancon | Panama |
| *Oophaga pumilio* | F | FT | Bocas del Toro | Panama |
| *Atelopus varius* | F | CB | Gamboa Amphibian Recovery Center | Panama |
| *Atelopus varius* | F | CB | Gamboa Amphibian Recovery Center | Panama |
| *Rhinella icterica* | F | FT | São Paulo (SP) | Brazil |
| *Elachistocleis panamensis* | NR | FT | Gamboa | Panama |

Abbreviations: M=Male; F=Female; NR=Not recorded; CB=Captive breeding; FT=Field trip.

### B. Eye size data

| Species | Eye length (mm) | Source |
|---|---|---|
| *Bombina orientalis* | 4.1 | MZUSP 115014, 59118, 59119 |
| *Brachycephalus ephippium* | 1.7 | Thais Condez* |
| *Ischnocnema parva* | 2.0 | Mariane Targino* |
| *Craugastor fitzingeri* | 5.8 | [1] |
| *Thoropa taophora* | 6.3 | Ariadne F. Sabbag* |
| *Hylodes asper* | 5.4 | [2] |
| *Hylodes phyllodes* | 3.8 | [2] |
| *Hylodes pipilans* | 3.1 | [4] |
| *Agalychnis callidryas* | 5.1 | [6] |
| *Pithecopus hypochondrialis* | 4.5 | [7] |
| *Bokermannohyla astartea* | 4.3 | [9] |
| *Bokermannohyla hylax* | 6.4 | MZUSP 71293, 153653, 153655, 98115 |
| *Boana geographica* | 4.4 | [12] |
| *Boana rosenbergi* | 7.6 | [13,14] |
| *Scinax ruber* | 3.2 | [16] |

| Species | Eye length (mm) | Source |
|---|---|---|
| *Xenohyla truncata* | 3 | [18] |
| *Dendropsophus ebraccatus* | 3.1 | MZUSP 101425 |
| *Dendropsophus microcephalus* | 2.7 | MZUSP 126217, 126218, 92538 |
| *Engystomops pustulosus* | 2.6 | MZUSP 135118, 135119, 5606 |
| *Physalaemus cuvieri* | 2.9 | MZUSP 151705, 151698, 151713, 151717, 151704, 151693, 151680, 151696, 151701, 151697, 151688, 151679, 151689 |
| *Leptodactylus insularum* | 3.5 | MZUSP 5454, 143068 |
| *Leptodactylus fragilis* | 3 | MZUSP 143063, 143064 |
| *Leptodactylus tapiti* | 2.3 | MZUSP 74314, 74315, 74325, 74326, 74327, 74328, 74329, 74332 |
| *Teratohyla spinosa* | 3.2 | Marco A. Rada* |
| *Cochranella granulosa* | 2.8 | Roberto Ibáñez |
| *Allobates femoralis* | 3.2 | [3] |
| *Aldephobates castaneoticus* | 2.6 | [5] |
| *Dendrobates auratus* | 2.8 | MZUSP 100169, 100170, 100171, 100172, 100173, 100174, 100175, 100176 |
| *Oophaga pumilio* | 2.2 | [8] |
| *Atelopus varius* | 3.3 | [10] |
| *Rhinella ornata* | 6.4 | [11] |
| *Rhinella icterica* | 11.5 | MZUSP 123383, 123386, 123536, 123543 |
| *Bufo bufo* | 5 | [15] |
| *Elachistocleis panamensis* | 1.7 | [17] |
| *Lithobates catesbeianus* | 8.2 | [19] |
| *Lithobates pipiens* | 6.3 | [20] |
| *Rana temporaria* | 4.4 | [21] |

(*) Unpublished data provided by the authors. MZUSP: Zoology Museum of University of São Paulo (Brazil). The numbers indicate the specimen IDs in the Herpetology collection.

### C. Comparison of different eye size measuring methods

To validate the use of eye sizes available in the taxonomic literature (i.e. measured as the distance between anterior and posterior edge of the eye in intact specimens), we compared the values that we collected with those reported for the axial diameter of enucleated eyes in the species for which they were available [22]. The linear regression shows an excellent fit, demonstrating that eye sizes as reported in species' descriptions can be used as a valid proxy for actual axial lengths.

| Species | Eye size, intact specimens (mm) | Axial length, enucleated eye (mm) |
|---|---|---|
| *Bombina orientalis* | 4.1 | 4.3 |
| *Brachycephalus ephippium* | 1.7 | 2.2 |
| *Boana rosenbergi* | 7.6 | 8 |
| *Atelopus varius* | 3.3 | 3.3 |
| *Dendrobates auratus* | 2.8 | 3.1 |
| *Oophaga pumilio* | 2.2 | 2.6 |
| *Teratohyla spinosa* | 2.8 | 3.0 |
| *Engystomops pustulosus* | 2.6 | 3.1 |
| *Physalaemus cuvieri* | 2.9 | 3.2 |
| *Elachistocleis panamensis* | 1.7 | 1.8 |

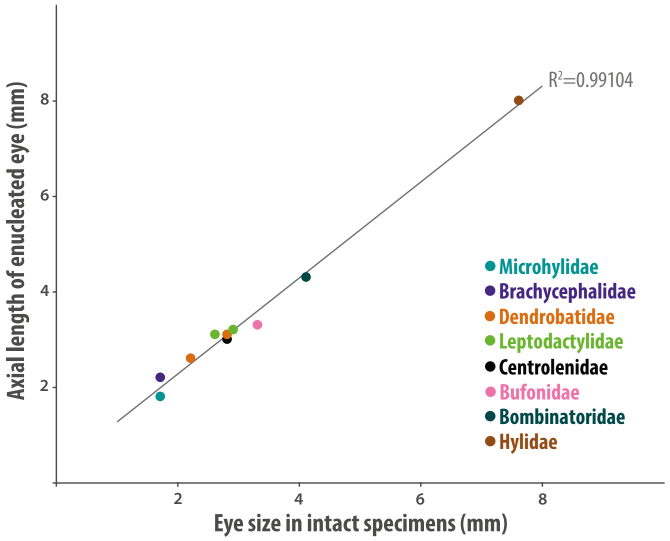

**D. Pupil shape scoring**

For pupil shape, we visually inspected photographs of each of the species available online and scored each of them on a dichotomic scale. The two scores were defined as “round” for those pupils that are either completely circular or just slightly oval/elliptical (and thus cover the periphery of the lens when contracting), and “elongate” for those that are clearly oval or slit-shaped (and thus contract proportionately more along one axis, while in the perpendicular axis most of the lens remains exposed to light). The threshold for “slightly oval” vs. “clearly oval” was set at:

$$\frac{Vertical\ pupil\ length}{Horizontal\ pupil\ length} = 0.75$$

**E. Species replacement in the phylogenetic tree**

Seven of the species in our dataset were absent in the tree from Pyron [23], thus we accommodated them as follows:

(1) We replaced *Rhinella crucifer* with *R. ornata*, which was a synonym of *R. crucifer* until it was resurrected by Baldissera and coworkers [11], and the two species are phylogenetically proximate in the *R. crucifer* species group [24].

(2) We replaced *Leptodactylus longirostris* with *L. fragilis*, both of which are in the same clade of three species in the *L. fuscus* species group [25].

(3) We replaced *L. bolivianus* with its sister species in the *L. latrans* species group, *L. insularum* [25].

(4) We replaced *L. gracilis* with *L. tapiti*; among the species in Pyron's tree [23], *L. tapiti* is phylogenetically closest to *L. gracilis* in the *L. fuscus* species group [25].

(5) We replaced *Hylodes dactylocinus* with the closely related species *H. asper* ([26]; R. Montesinos and T. Grant, unpublished data).

(6) Given their resemblance and close relationship ([4]; R. Montesinos and TG, unpublished data), we inserted *H. pipilans* as sister to *H. phyllodes* with branch lengths of zero and used the original *H. phyllodes* branch length as the branch length for the inclusive clade.

(7) Finally, we replaced *Elachistocleis bicolor* with *E. panamensis*, which is the sister species of the *E. bicolor* + *E. ovalis* clade [27].

**F. Phylogenetic tree used for statistical analyses**

Modified from [23] to include only the species included in the present study (see main text and S1E, above).

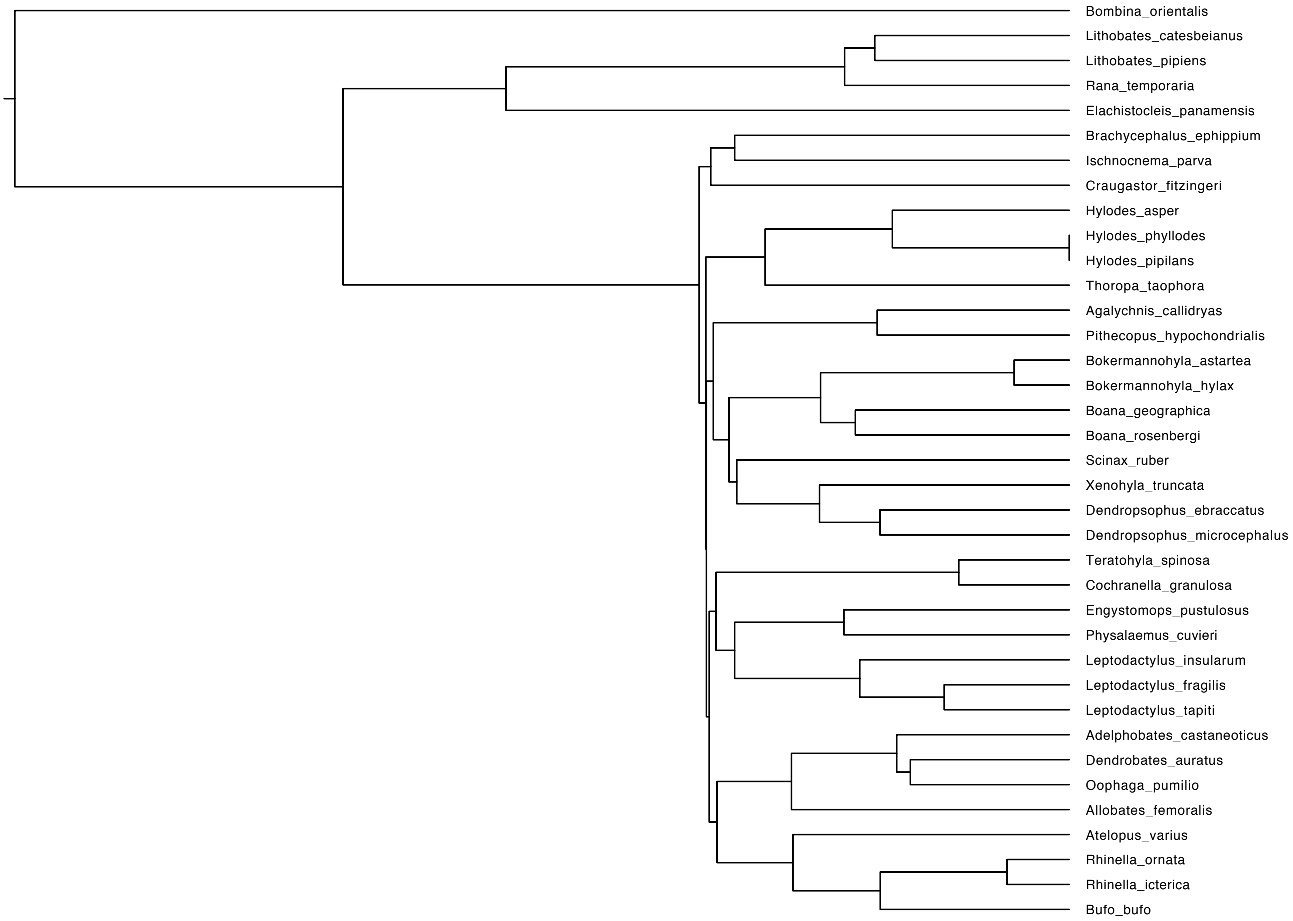

### G. Corneal transmittance

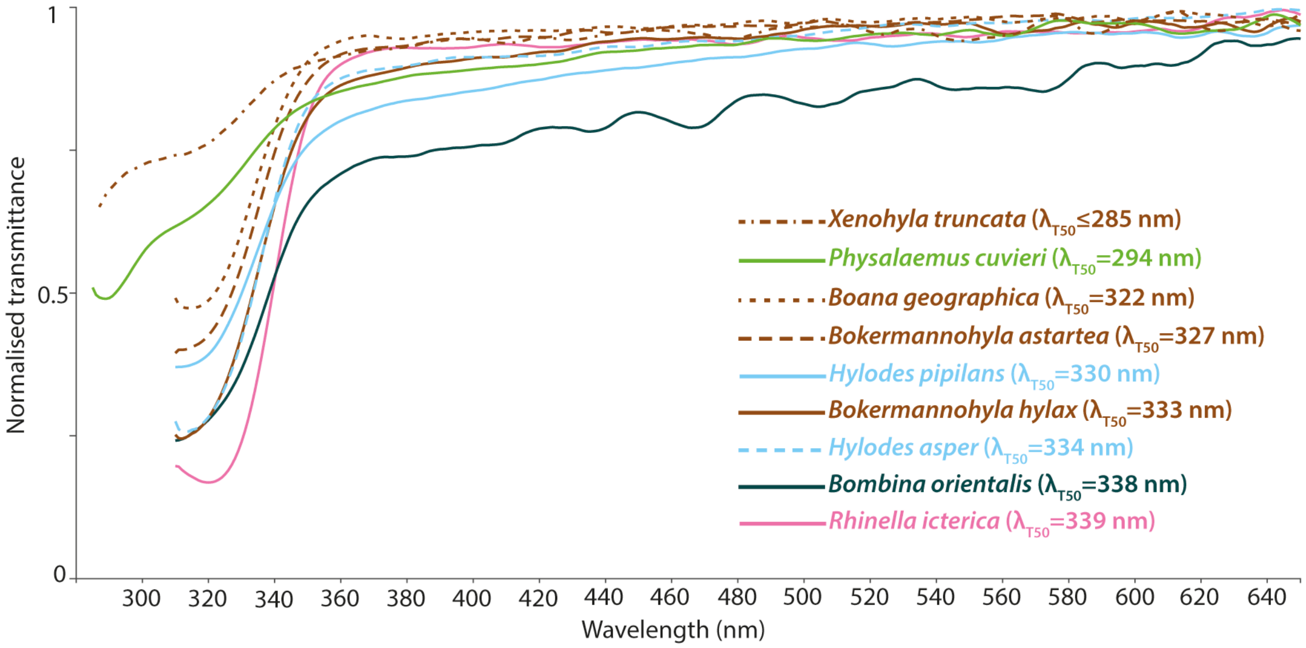

### H. Lens transmittance curves of species not shown in Figure 3 of the main text

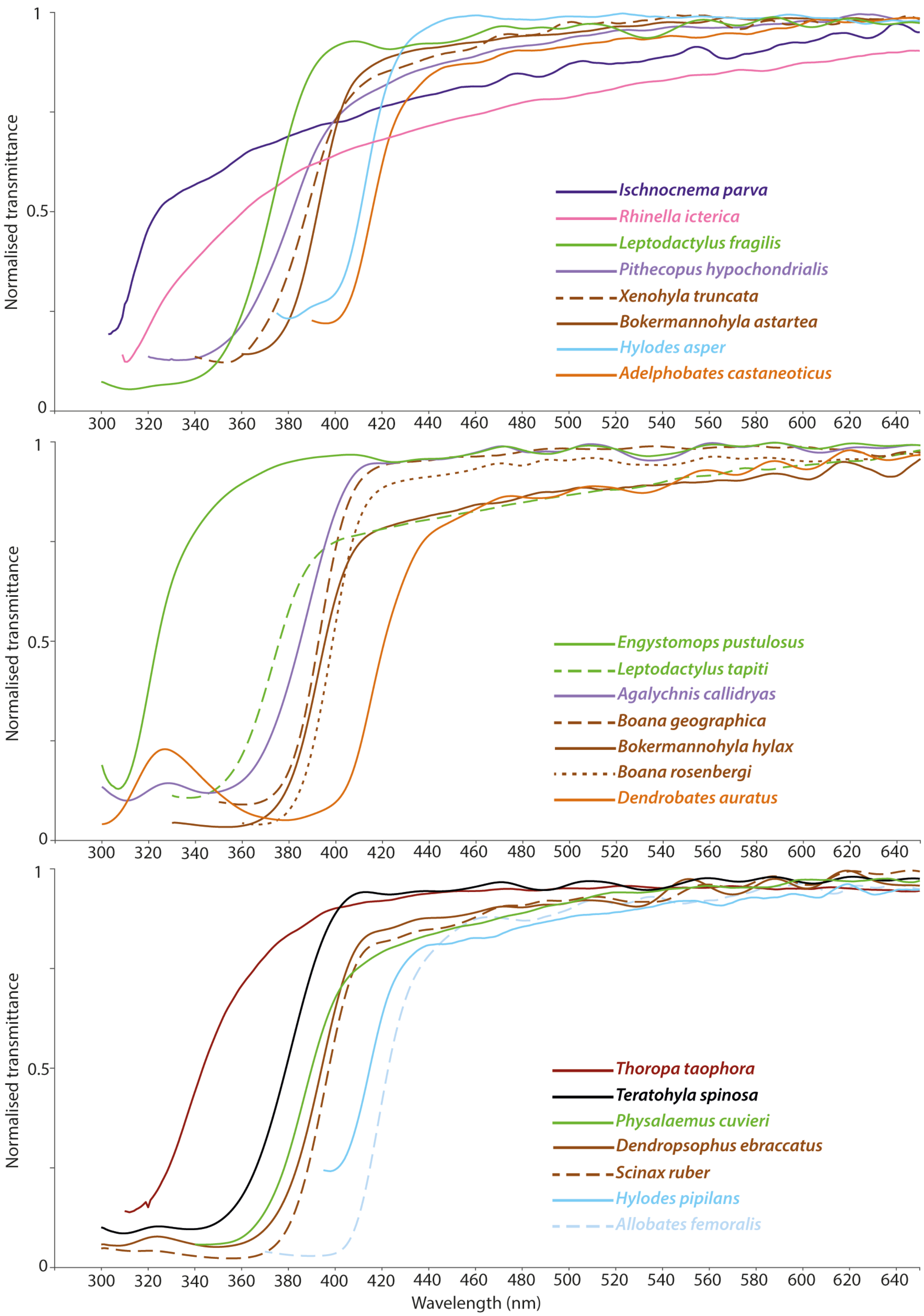

### I. Statistical analyses

Results (regression coefficients, b, and corresponding p-values) of phylogenetic generalized linear mixed model analysis of binary data using the model Lens $\lambda_{T50}$ ~ Eye size, excluding *Bombina orientalis*.

| Variable | b | p |
|---|---|---|
| Eye size ($\lambda$ = 0) | 0.416 | 0.8664 |
| Eye size ($\lambda$ = 1) | 0.0137215 | 0.2498 |

Results (regression coefficients, b, and corresponding p-values) of phylogenetic generalized linear mixed model analysis of binary data using the model Diel pattern ~ Lens $\lambda_{T50}$ + Eye size + Pupil shape, including *Bombina orientalis*.

| Variable | b | p |
|---|---|---|
| Lens $\lambda_{T50}$ | 0.031269 | 0.1640 |
| Eye size | -0.568173 | 0.3113 |
| Pupil shape | -1.219676 | 0.4019 |

Results (regression coefficients, b, and corresponding p-values) of phylogenetic generalized least squares analysis of continuous data using the model Elevation ~ Lens $\lambda_{T50}$ + Eye size + Pupil shape, including *Bombina orientalis*.

| Variable | Pagel's $\lambda$ = 0 | | Pagel's $\lambda$ = 0.1 | | Pagel's $\lambda$ = 0.5 | | Pagel's $\lambda$ = 1 | |
|---|---|---|---|---|---|---|---|---|
| | b | p | b | p | b | p | b | p |
| Lens $\lambda_{T50}$ | -0.4589 | 0.8399 | -0.4646 | 0.8383 | -0.45261 | 0.8624 | 1.43842 | 0.6183 |
| Eye size | 26.0041 | 0.5208 | 26.0506 | 0.5204 | 25.58508 | 0.5454 | 5.38262 | 0.9089 |
| Pupil shape | -22.9854 | 0.9012 | -23.2553 | 0.9001 | -43.06161 | 0.8249 | -70.46774 | 0.7386 |

Results (regression coefficients, b, and corresponding p-values) of phylogenetic generalized least squares analysis of continuous data using the model Latitude ~ Lens $\lambda_{T50}$ + Eye size + Pupil shape, including *Bombina orientalis*.

| Variable | Pagel's $\lambda$ = 0 | | Pagel's $\lambda$ = 0.1 | | Pagel's $\lambda$ = 0.5 | | Pagel's $\lambda$ = 1 | |
|---|---|---|---|---|---|---|---|---|
| | b | p | b | p | b | p | b | p |
| Lens $\lambda_{T50}$ | -0.10722 | 0.1223 | -0.10626 | 0.1253 | -0.07397 | 0.2913 | 0.071184 | 0.2670 |
| Eye size | 1.60505 | 0.1909 | 1.58655 | 0.1949 | 0.87073 | 0.4402 | -1.752291 | 0.0981 |
| Pupil shape | 3.97034 | 0.4766 | 3.95965 | 0.4765 | 3.20465 | 0.5371 | 3.063991 | 0.5116 |

Results (regression coefficients, b, and corresponding p-values) of phylogenetic generalized linear mixed model analysis of binary data using the model Diel pattern ~ Lens $\lambda_{T50}$ + Eye size + Pupil shape, excluding *Bombina orientalis*.

| Variable | b | p |
|---|---|---|
| Lens $\lambda_{T50}$ | 0.031062 | 0.1710 |
| Eye size | -0.567791 | 0.3124 |
| Pupil shape | -1.219609 | 0.4031 |

Results (regression coefficients, b, and corresponding p-values) of phylogenetic generalized least squares analysis of continuous data using the model Elevation ~ Lens $\lambda_{T50}$ + Eye size + Pupil shape, excluding *Bombina orientalis*.

| Variable | Pagel's λ = 0 | | Pagel's λ = 0.1 | | Pagel's λ = 0.5 | | Pagel's λ = 1 | |
|---|---|---|---|---|---|---|---|---|
| | b | p | b | p | b | p | b | p |
| Lens $\lambda_{T50}$ | -1.3779 | 0.5995 | -1.3698 | 0.6023 | -0.7672 | 0.7950 | 1.8885 | 0.5730 |
| Eye size | 25.3422 | 0.5346 | 25.4029 | 0.5342 | 24.9619 | 0.5646 | 9.6770 | 0.8420 |
| Pupil shape | -12.3170 | 0.9475 | 12.8484 | 0.9453 | -43.7151 | 0.8279 | -13.3644 | 0.9520 |

Results (regression coefficients, b, and corresponding p-values) of phylogenetic generalized least squares analysis of continuous data using the model Latitude ~ Lens $\lambda_{T50}$ + Eye size + Pupil shape, excluding *Bombina orientalis*.

| Variable | Pagel's λ = 0 | | Pagel's λ = 0.1 | | Pagel's λ = 0.5 | | Pagel's λ = 1 | |
|---|---|---|---|---|---|---|---|---|
| | b | p | b | p | b | p | b | p |
| Lens $\lambda_{T50}$ | -0.023462 | 0.7481 | -0.023541 | 0.7469 | -0.022762 | 0.7583 | 0.0766823 | 0.2623 |
| Eye size | 1.665369 | 0.1488 | 1.639267 | 0.1542 | 0.741399 | 0.4951 | -1.6569201 | 0.0992 |
| Pupil shape | 2.998050 | 0.5667 | 2.994924 | 0.5661 | 2.345081 | 0.6419 | 1.5587347 | 0.7291 |

### J. Lens transmittance vs. eye size in a subset of anuran species with putatively unpigmented lenses

Each data point represents one species, and the data point with a black outline was obtained from a previous study [28].

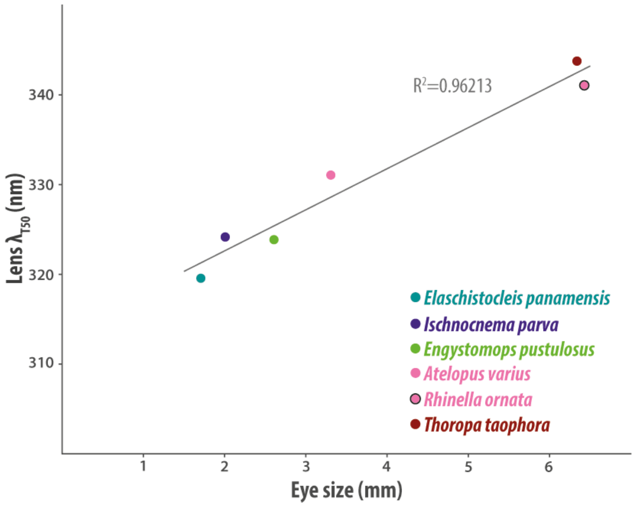
